# Quantitative mathematical modeling of clinical brain metastasis dynamics in non-small cell lung cancer

**DOI:** 10.1101/448282

**Authors:** M. Bilous, C. Serdjebi, A. Boyer, P. Tomasini, C. Pouypoudat, D. Barbolosi, F. Barlesi, F. Chomy, S. Benzekry

**Affiliations:** MONC team, Inria Bordeaux Sud-Ouest, Talence, France.; Institut de Mathématiques de Bordeaux, Bordeaux University, Talence, France.; SMARTc Unit, Center for Research on Cancer of Marseille (CRCM), Inserm UMR 1068, CNRS UMR 7258, Aix-Marseille University U105. Marseille, France.; CRCM, Inserm UMR 1068, CNRS UMR 7258, Aix Marseille University, Assistance Publique Hôpitaux de Marseille. Marseille. France.; Radiation oncology department, Haut-Lévêque Hospital, Pessac, France; Clinical oncology department, Institut Bergonié, Bordeaux, France

## Abstract

Brain metastases (BMs) are associated with poor prognosis in non-small cell lung cancer (NSCLC), but are only visible when large enough. Therapeutic decisions such as whole brain radiation therapy would benefit from patient-specific predictions of radiologically undetectable BMs. Here, we propose a mathematical modeling approach and use it to analyze clinical data of BM from NSCLC.

Primary tumor growth was best described by a gompertzian model for the pre-diagnosis history, followed by a tumor growth inhibition model during treatment. Growth parameters were estimated only from the size at diagnosis and histology, but predicted plausible individual estimates of the tumor age (2.1-5.3 years). Multiple metastatic models were assessed from fitting either literature data of BM probability (n = 183 patients) or longitudinal measurements of visible BMs in two patients. Among the tested models, the one featuring dormancy was best able to describe the data. It predicted latency phases of 4.4 - 5.7 months and onset of BMs 14 - 19 months before diagnosis. This quantitative model paves the way for a computational tool of potential help during therapeutic management.

## Introduction

Lung cancer is the leading cause of cancer-related death worldwide^1^. Nearly 80% of lung cancers are non-small cell lung cancer (NSCLC) and 60% of them are diagnosed at the metastatic stage^2^. Brain metastases (BMs) affect more than 20% of patients with NSCLC^3,4^. Despite recent advances in this field, BMs remain a major challenge as they are associated with poor prognosis^5–8^. In addition, BMs are responsible for disabling symptoms, with impaired patients’ reported outcomes.

Several biological and clinical open questions can be formulated: 1) What was the pre-diagnosis course of a patient presenting with NSCLC? 2) For a patient developing overt BMs, when did cerebral invasion first occur? 3) Were most of the BMs spread by the primary tumor (PT) or did BMs themselves represent a significant source of secondary spread^9–11^? 4) How did growth kinetics compare between the PT and the BMs? Were there dormancy periods during the BMs history^12,13^? 5) For a patient with no or few (≤ 5) BMs at a given time point, what is the risk and extent of the occult disseminated disease in the brain (if any)?

Answers to these questions would have direct clinical utility. The clinical follow-up and planning of cerebral magnetic resonance imaging (MRI) could indeed highly benefit from individualized predictions of the probability of BM relapse or development. Moreover, the prediction of the risk and timing of BM may help clinical decision between localized (e.g. stereotactic radiotherapy), pan-cerebral (whole-brain radiotherapy (WBRT)) and/or systemic treatment. As of today, prophylactic cranial irradiation in the management of NSCLC is not recommended despite a reduction in the risk of BMs, due to the lack of overall survival benefit and important neuro-cognitive toxicities^6,14,15^. Despite recent improvement in radiotherapy techniques, with the development of hypocampal-sparing WBRT^16^, whole brain volume reduction and neurocognitive function decline were described^17^. This highlights the need for a better prediction of patients with highest risk of BM development or relapse, in order to select patients with the higher benefit/risk ratio for WBRT. Several phase III trials were conducted but no firm conclusion applicable for the entire patient population could be drawn^18,19^. The prediction of the risk of BM development and indication for WBRT to prevent brain dissemination is particularly interesting for specific populations such as *EGFR* or *ALK* positive NSCLC patients, for whose incidence of BMs is high^20–22^. This points to the need of rational tools to decide therapeutic action in a patient-specific way.

For such aims, quantitative mathematical modeling may be of considerable help, by inferring underlying processes from observed data or providing useful numerical tools in the era of personalized medicine^23,24^. However, despite numerous studies in mathematical and computational oncology, the majority of the efforts have to date focused on mathematical models at the scale of a single tumor^25^. Moreover, the comparison of the models to empirical data remains infrequent. Historically, modeling efforts in the field of metastasis were first initiated by statistical models phenomenologically describing relapse hazards^26–29^. In the 1970’s, Liotta et al. pioneered the development of biologically-based and experimentally-validated models for all the main steps of the metastatic process^30^. Since then, only a small number of studies addressed this topic^31–40^. Of specific relevance to the current work, the Iwata model^31^ introduced a size-structured population approach to capture the time development of a population of regional metastases from a hepatocellular carcinoma. It was further extended and studied from the mathematical and numerical viewpoints^41–43^, and was found able to reproduce experimental *in vivo* data^37–39^.

To understand biological dynamics of metastasis, animal studies allowing to capture the natural course of the disease^37,38^ and the effect of therapeutic interventions in controlled environments^39^ provide valuable data for quantitative analysis, although with limited clinical relevance. Experimental procedures are tedious and often only provide access to the total metastatic burden, neglecting its distribution into distinct metastatic lesions when no anatomical imaging modalities are employed^38,39^.

Individual-level clinical data of metastatic development are also challenging to obtain because precise number and size of existing lesions are not routinely collected. Apart from the landmark work of Iwata et al.^31^ (which only included one untreated patient with hepatocellular carcinoma), no study has yet provided a quantitative modeling analysis of longitudinal data of individual-level number and size of metastases. The purpose of the current work is to develop this approach for BMs dynamics in NSCLC, in order to address the biological and clinical questions stated above.

## Results

### Pre-diagnosis natural course of lung primary tumors

We investigated two possible growth models for the natural course of lung PTs: exponential and Gompertz. Exponential growth is the simplest model expressing uncontrolled proliferation and is often well adapted to describe tumor growth kinetics during limited observation periods^44^. However, it has been shown that on longer timeframes (typically under volumes increases of 100 to 1000-fold), the specific growth rate of tumors decreases^45^, which is well captured by the Gompertz model^45–47^. For calibration of the parameters, we used the data of the PT size and mean doubling time at diagnosis (see Methods). The latter was assumed to be histology-dependent and retrieved from a meta-analysis comprising a total of 388 adenocarcinomas and 377 squamous cell carcinomas^44,48^ (see supplementary Table S1). For the Gompertz model (two parameters), we additionally assumed a carrying capacity of 10^12^ cells^49^. Under such assumptions, we obtained predictions of pre-diagnosis phases of 19 years versus 5.4 years in the exponential and Gompertz cases, respectively, for an adenocarcinoma with diameter 35 mm (median value from ref.^50^), see Figure S1. The first prediction seems highly unrealistic compared with previous reports estimating the age of lung primary tumors to be 3-4 years old^51^, based on a different method using time to recurrence^52^. The Gompertz prediction on the other hand is rather consistent with the literature range. Moreover, the obtained value of the cellular proliferation rate was realistic, corresponding to a length of the cell cycle 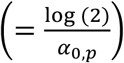 of 24.4 hours^53^.

We therefore concluded that the Gompertz model was best adapted to describe the pre-diagnosis natural history of the PT.

### Probability of brain metastasis as a function of primary tumor size can be described by a mechanistic computational model of metastasis

PT size was reported by Mujoomdar et al.^50^ as a major predictive factor of BM in NSCLC. We investigated whether our mechanistic model of BM was able to fit the data from these authors, at the population level (see Methods for the definition of the model). To do so, we assumed a statistical distribution associated with the inter-individual variability for the metastatic dissemination parameter *μ* (see Methods). The growth parameters were determined as above, i.e. from histology only. This approach yielded size-dependent population probabilities of BM occurrence, which could be compared to the data. The model fits were reasonably accurate, for both histological types (Figure 1). Importantly, this demonstrates the descriptive power of our model, which is based on mechanistic simulations of the underlying biological process. In this regard, this is superior to a mere fit of a biologically agnostic statistical model such as a linear regression. Notably, this result was obtained with a minimal number of parameters to describe the inter-patient heterogeneity in metastatic potential. Namely, only two parameters defining the lognormal statistical distributions of *μ* in the populations. The coefficients of variation were very large (> 6,000%). The median value *μ*_pop_ gives a quantitative way to measure median BM aggressiveness. It was found two log_10_ orders of magnitude larger for adenocarcinomas than for squamous cell carcinomas (*μ*_pop_ = 1.39 × 10^−11^ vs *μ*_pop_ = 1.76 × 10^−13^).

**Figure 1:**
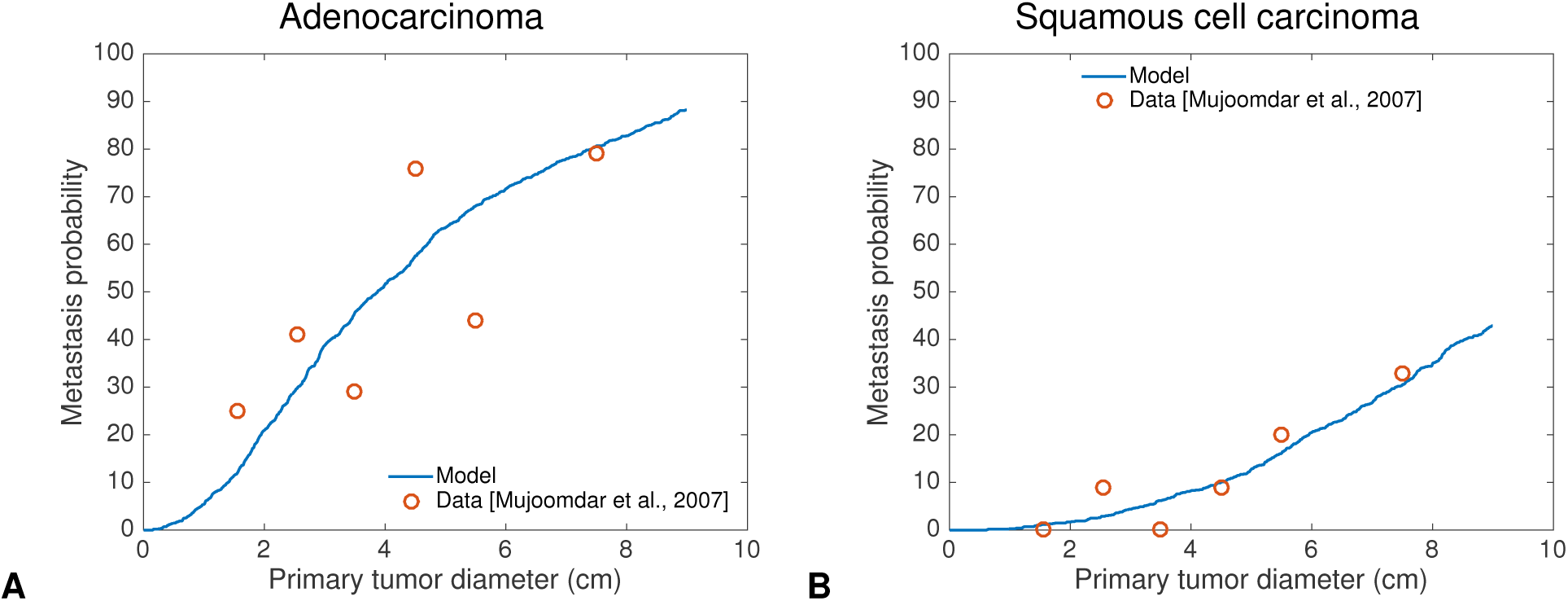
Probability of brain metastasis as a function of the primary tumor size. Fit of the mechanistic model for probability of BM to data from Mujoomdar et al. A. Adenocarcinoma data (n=136). Inferred values for the distribution of *µ*: *µ*_pop_ = 1.39 × 10^−11^ cell^−1^· day^−1^ and *µ*_σ_ = 1.24 × 10^−9^ cell^−1^ · day^−1^ B. Squamous cell carcinoma data (n=47). Inferred values for the distribution of *µ*: *µ*_pop_ = 1.76 × 10^−13^ cell^−1^· day^−1^ and *µ*_σ_ = 1.11 × 10^−11^ cell^−1^ · day^−1^

### Model-based comparison of biological scenarios with individual longitudinal data suggests dormancy

#### Data

These consisted of 10 and 11 PT size measurements, and 47 and 16 BM size measurements in two patients, respectively (one with an adenocarcinoma, the other with a squamous cell carcinoma). We first analyzed the adenocarcinoma patient (Figure 2) to develop the model and used the second patient to investigate the generalizability of our findings. In the adenocarcinoma patient, the PT first responded to systemic therapy before slowly regrowing (Figure 2A). A first distant BM was detected 20 months after diagnosis, and kept growing uncontrolled afterwards (Figure 2B). Other BMs appeared during follow-up (Figure 2C), reaching a total of 20 BMs at 48 months, date of last examination (Figure 2D).

**Figure 2:**
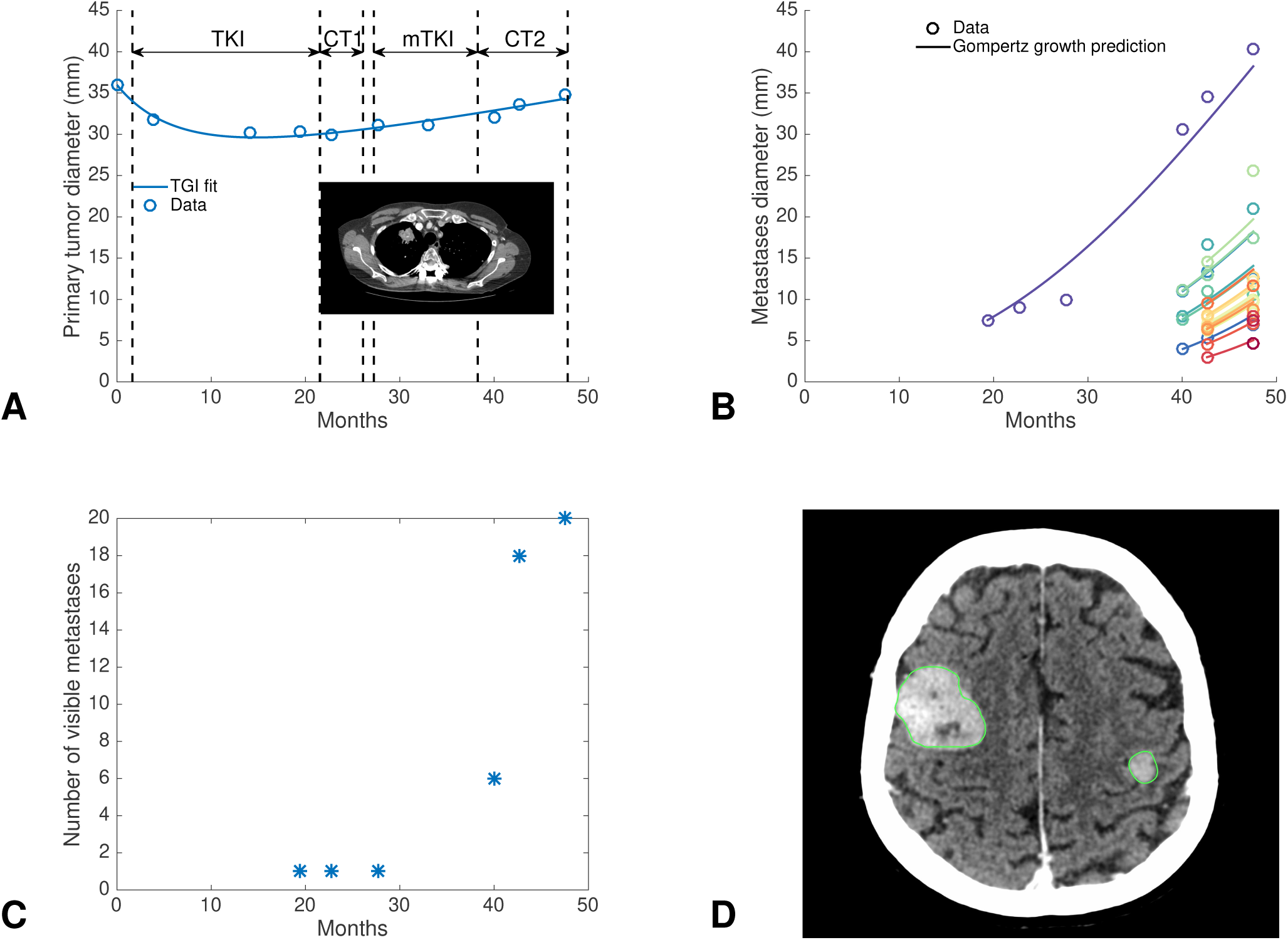
Data of primary tumor and brain metastases in a patient with non-small cell lung cancer. A. Post-diagnosis kinetics of the primary tumor largest diameter, measured on follow-up computerized tomography images (inlet). This EGFR mutated patient was treated first with a tyrosine kinase inhibitor (erlotinib) then with several rounds of additional systemic treatments upon relapse (cytotoxic chemotherapy (docetaxel), re-challenge with erlotinib and a second cytotoxic agent (gemcitabine)), as indicated by arrows and dashed lines in the figure. The solid line corresponds to the model fit during treatment (tumor growth inhibition (TGI) model). B. Growth kinetics of the brain metastases. The solid line corresponds to Gompertz growth predictions based on parameters estimated from the primary tumor size at diagnosis and histological type, using as initial condition the first observation of each BM. C. Time course of the apparition of visible metastases. D. Cerebral magnetic resonance image showing two brain metastases 48 months post-diagnosis (other brain lesions not visible on this slice). *EGFR = Epithelial Growth Factor Receptor, TKI = Tyrosine Kinase Inhibitor, mTKI = maintenance Tyrosine Kinase Inhibitor, CT = (cytotoxic) chemotherapy. Time is in months from diagnosis*.

#### Growth kinetics

To model the effect of systemic treatment on the PT, we found that a tumor growth inhibition model^54^ (equation (1)) was able to adequately fit the data (Figure 2A).

Interestingly, for the BM kinetics, we found that the untreated PT Gompertz model predicted surprisingly well the observed data (Figures 2B and S2). Specifically, when employing the Gompertz parameters used for the pre-diagnosis PT history, and using initial conditions on each BM, the future growth curves were accurately predicted. This indicated that: 1) for this patient, the BMs did not respond to systemic therapy, at least from the time of first observation, 2) it is reasonable to assume that all BMs grew at the same growth rate, 3) the BMs growth rate might be similar to the PT growth rate, at least during the clinically overt phase.

#### Quantitative assessment of five biologically-based models of metastatic dissemination and colonization

However, this mere description of the BMs growth is not satisfactory as a model of systemic disease, since the dissemination part is absent. In particular, no model is given for the initiation times of the BMs. To include the dissemination component of the metastatic process, we relied on a modeling framework first initiated by Iwata et al.^31^ (see Methods).

Five models, corresponding to five possible biological scenarios, were investigated (Figure 3). These included: (A) a “basic” model with same growth model and parameters for the PT and the BMs and no secondary dissemination (i.e. BMs spread only by the PT); (B) a model allowing different growth parameters for the PT and the BMs; (C) a model with secondary dissemination, i.e. the ability of BMs to spread BMs themselves^31^; (D) a model allowing a delay before initiation of metastatic ability of the PT (the so-called linear model in which dissemination occurs at a late stage, opposed to the parallel model where dissemination is an early event^49^) and (E) a model allowing BM dormancy phases, i.e. time periods with null growth rate^12,13,55^. Each of these scenario was translated into corresponding mathematical equations (see Methods).

**Figure 3:**
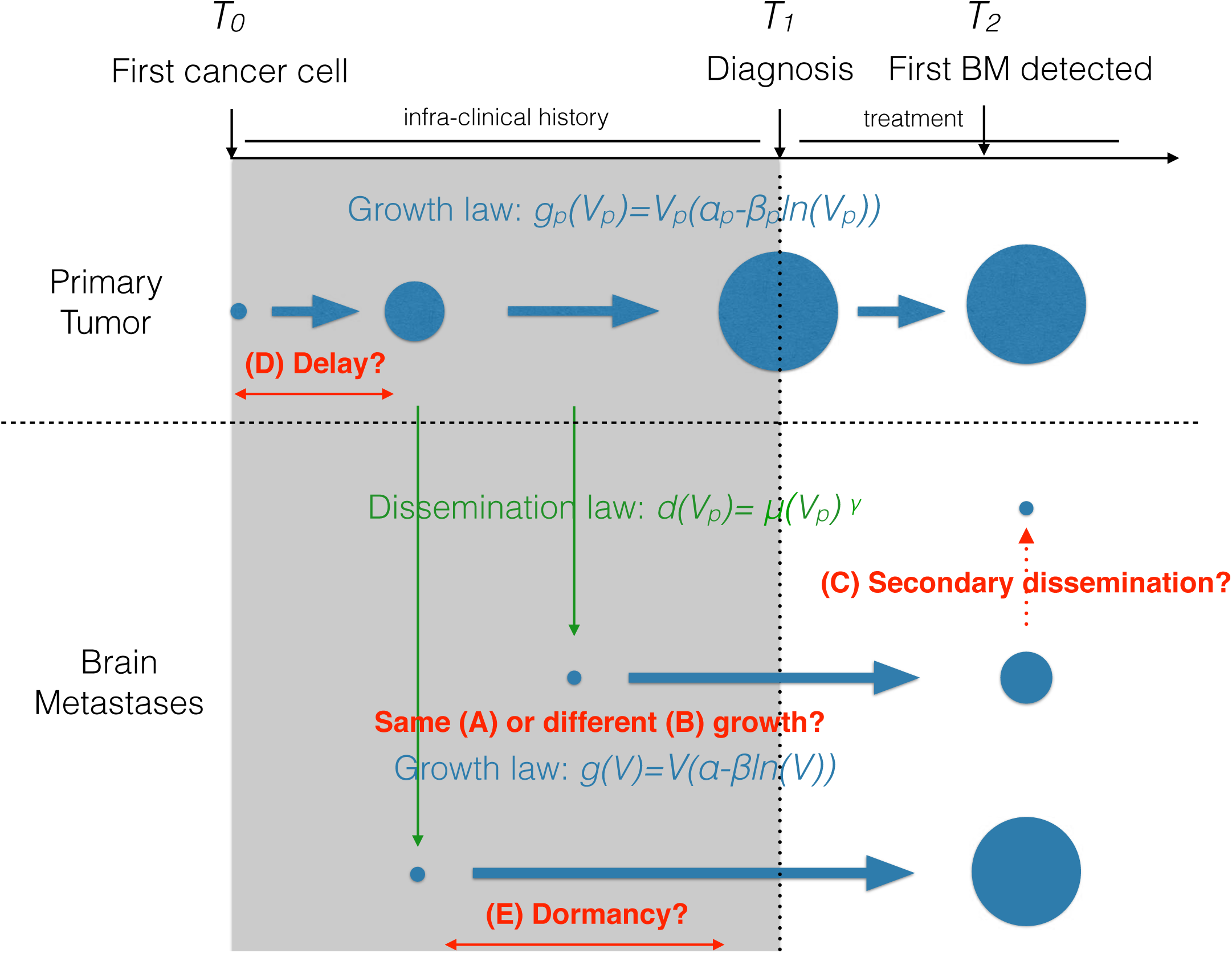
Schematic of the models. Several biological hypotheses about metastatic dynamics were formalized as mathematical models and tested against the data. These included: same (A) or different (B) growth rates between primary and secondary tumors; only primary dissemination from the primary tumor or (C) secondary dissemination from brain metastases themselves; (D) possibility of a delay between cancer initiation and onset of metastatic ability; and (E) the possibility of a dormancy lag time between metastatic spread and growth initiation.

The models exhibited different descriptive power, as quantified by the best-fit value of the objective function (Table 1). The best description of the data was obtained by the dormancy model (Figure 4). Importantly, this model achieved accurate goodness-of-fit for 47 BM size measurements over 6 time points (Figure 4A-B) with only 3 parameters fitted from the BM sizes data (*μ*, *τ* and *γ*). Consequently, parameter identifiability was found excellent (relative standard errors < 5%, Table 2). Apparition of the first BM above the visibility threshold was also accurately described (19 vs 20 months, model vs data, Figure 4C). The model overestimated the number of small visible BMs at 40 months, possibly because a lot of these were present but too small to be visible at MRI. Indeed, a lot of BMs were present at 43 months – in accordance with the model (Figure 4A) – which thus emphasizes its predictive potential.

**Figure 4:**
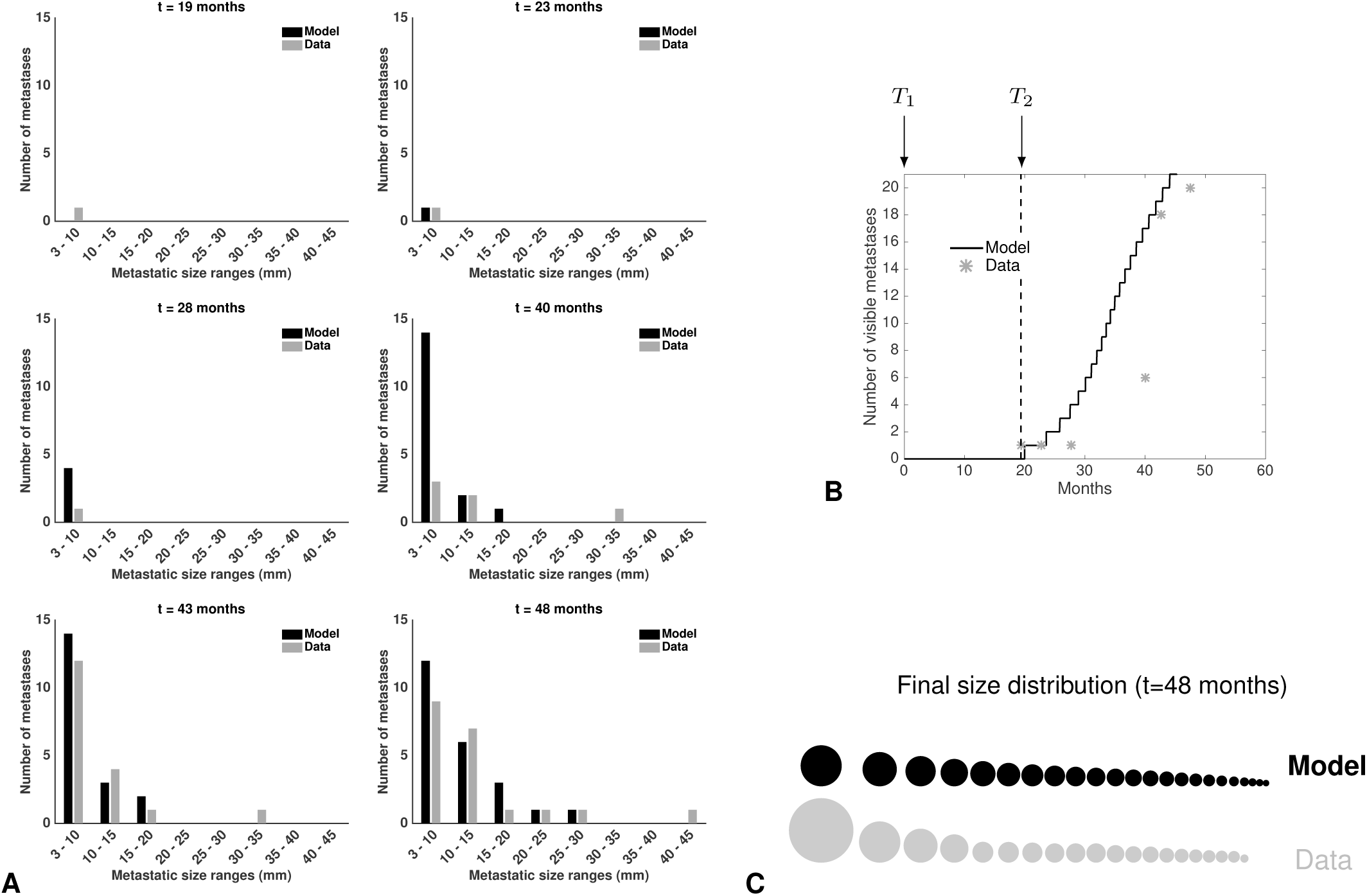
Fit of the dormancy model. A. Time course of the visible brain metastases (BMs) size distributions during follow-up. Comparison between model calibration and data. B. Time course of the number of visible BMs. T1 = time of diagnosis. T2 = detection time of the first brain metastasis. C. Comparison of the BM size distribution between the model fit and the data at last follow-up (48 months post-diagnosis). *Time is in months from diagnosis*.

**Table 1:** Minimal value of the objective function obtained when fitting each of the models to the data

| Model | Patient 1 | Patient 2 |
| --- | --- | --- |
| Basic | 5.51 | 2.53 |
| Secondary | 5.43 | 2.3 |
| Delay | 5.23 | 1.53 |
| Dormancy | 4.93 | 1.71 |
| Diff. growth | 4.95 | 1.79 |

The objective function was the sum of squared residuals (see expression (7)).

**Table 2:** Patient-specific parameters

| Parameter | Meaning | Unit | Patient 1 | Patient 2 | Determination |
| --- | --- | --- | --- | --- | --- |
| $S_d$ | PT size at diagnosis | mm | 36.0 | 53.7 | Data |
| Histology |  |  | Adenocarcinoma | Squamous cell carcinoma | Data |
| $T_d$ | Age of the PT at diagnosis | year | 5.3 | 2.1 | Predicted |
| Visibility threshold |  | mm | 3 | 3 | Fixed |
| $S_{0,p}$ | Initial size of the PT | cell | 1 | 1 | Fixed |
| $S_0$ | Initial size of a BM | cell | 1 | 1 | Fixed |
| $\alpha_{0,p}$ | Proliferation rate at one cell (PT) | day<sup>-1</sup> | 0.0284 | 0.0858 | Computed |
| $\beta_p$ | Exponential decrease of growth rate (PT) | day<sup>-1</sup> | $1.03 \times 10^{-3}$ | $3.10 \times 10^{-3}$ | Computed |
| $DT_{PT}$ | PT doubling time at 3 cm | day | 173 | 57 | Predicted |
| $\alpha_1$ | PT growth rate during relapse | day<sup>-1</sup> | $5.72 \times 10^{-4}$ (23.6) | 0.0101 ($5.28 \times 10^{-3}$) | Fit PT |
| $\kappa$ | PT log-kill effect | day<sup>-1</sup> | $4.46 \times 10^{-3}$ (20.6) | 0.0439 (1803) | Fit PT |
| $t_{res}$ | Half-life of treatment response | day | 149 ($1.05 \times 10^{-3}$) | 35.4 (0.998) | Fit PT |
| $\alpha_0$ | Proliferation rate at one cell (BM) | day<sup>-1</sup> | 0.0284 | 0.0858 | Computed |
| $\beta$ | Exponential decrease of growth rate (BM) | day<sup>-1</sup> | $1.03 \times 10^{-3}$ | $3.10 \times 10^{-3}$ | Computed |
| $\tau$ | Dormancy duration | day | 133 (4.23) | 171 (33.3) | Fit BM |
| $DT_{BM}$ | BM doubling time at 3 cm | day | 173 | 57 | Predicted |
| $\mu$ | Cellular metastatic potential | cell<sup>-1</sup> × day<sup>-1</sup> | $2.00 \times 10^{-12}$ (2.83) | $1.02 \times 10^{-12}$ (71.9) | Fit BM |
| $\gamma$ | Fractal (Hausdorff) dimension of the vasculature ($\times \frac{1}{3}$) | - | 1 | 1 | Fit BM* |
| $d$ | Dissemination rate at 3 cm | day<sup>-1</sup> | 0.0282 | 0.0144 | Predicted |

Values of the parameters from either the data (Data), the model assumptions (Fixed), estimated from direct computations from diagnosis data (Computed), fit of the TGI model to the PT data (Fit PT) or fit of the metastatic dormancy model to the brain metastases data (Fit BM), or computed from these estimations (Predicted). Values in parenthesis for fitted parameters correspond to the relative standard errors expressed in percent, i.e. $\frac{se}{est} \times 100$ where se is the standard error (square root of the covariance diagonal entry, see Materials and methods) and est is the parameter estimate. \* = for value of the parameter $\gamma$ , due to identifiability issues, only five possible values were considered during the fit procedure (see Methods).

The basic (A), secondary (C) and delay models (D) exhibited substantially worse goodness-of-fit (Table 1 and Figures S3-5), suggesting that their underlying hypotheses are insufficient to provide valid explanations of BM dynamics. In the delay model, the fit was improved compared to models (A) and (B) but the value of the delay *t*_d_ was found unrealistically large (4.8 years after the first cancer cell, corresponding to 6 months before diagnosis).

Finally, model (D) yielded a significant improvement of the goodness-of-fit for both the dynamics of the number of visible BMs and cumulative size distributions, while not deteriorating the practical identifiability of the parameters (Figure S6). Of note, the estimated values of the BM growth parameters were close to the PT ones. However, given the previous observation that during the clinical phase the BMs grew at a growth rate consistent with the one predicted for the PT, combined with the facts that the optimal objective value was achieved for the dormancy model and that it is more parsimonious (one parameter less), we selected the dormancy model for further analysis.

### Clinically relevant simulations of the disease course reveal times of BM initiation

From the quantitative calibration of the dormancy model to the data, several predictions of clinical interest can be made. The value of *γ* that generated the best-fit was 1, suggesting equal probability for all the cells within the PT to disseminate. In turn, this could indicate of a well-vascularized tumor, which might be prone to anti-angiogenic therapy. Interestingly, the value of *μ* inferred from this patient-level data was in the same range as the one inferred from the above population analysis (*μ* = 2 × 10^−12^ cell^−1^.day^−1^ versus *μ*_pop_ = 1.39 × 10^−11^ cell^−1^.day^−1^), giving further support to our approach.

Once calibrated from the data, our model allowed to simulate the predicted natural history of the disease. The supplementary movie S1 shows a simulation of the PT growth together with the apparition of the entire population of BMs (visible + invisible). Stars indicate tumors that are invisible (< 5 mm), and the BMs size distribution time course is also simulated. BMs represented in gray were born before the diagnosis, while BMs in white are the ones born after.

Apart from the age of the PT, prediction of birth times and invisible BM burden at any time point can be performed. Interestingly, we found that the first BM – which clinically appeared 19 months after diagnosis – was already present 14 months before clinical detection of the PT (Figure 5A). In fact, at the time of diagnosis, our model predicted the presence of 10 occult BMs, representing a total burden of 1,167 cells mostly distributed into the largest (first) BM (size 1,046 cells ≃ 0.126 mm), see Figure 5B. Notably, at the time the PT had reached the imaging detection limit, no metastasis had occurred yet. This suggests that if the disease had been detected earlier (e.g., by systematic screening), BM occurrence might have been prevented provided the tumor had been operable at this time. The amount of predicted BMs present at *T*_2_ was 28 tumors, with an overall BM burden of 1.7 × 10^7^ cells (Figure 5C). Therefore, provided that neuro-cognitive risks would be acceptable, the model would recommend WBRT or systemic treatment rather than localized intervention only. Together, these results demonstrate the potential clinical utility of the model for prediction of the invisible BM, in order to inform therapeutic decision.

**Figure 5:**
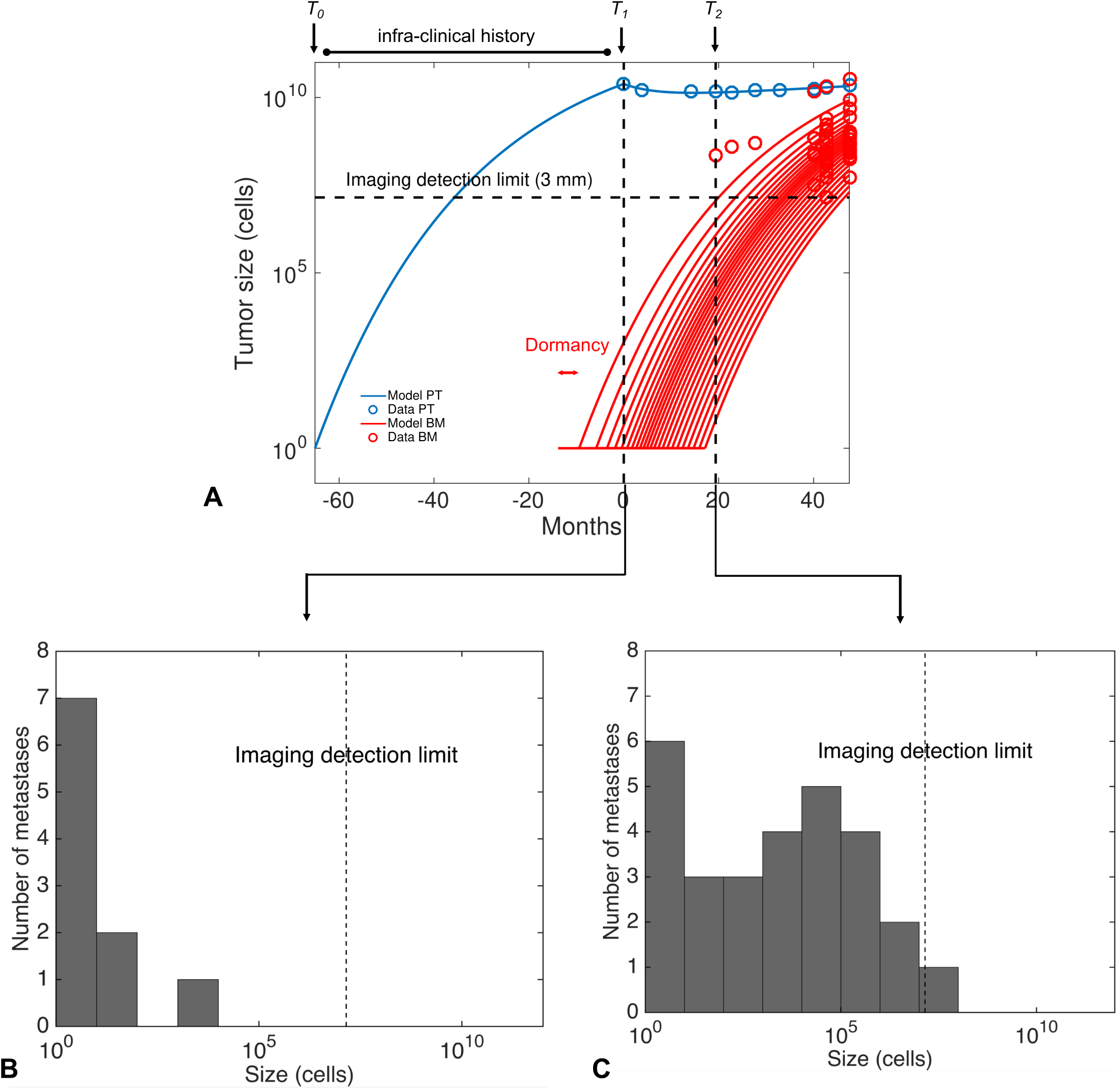
Model simulated predictions of the pre- and post-diagnosis time course of the primary and cerebral disease. A. Model-inferred growth kinetics of the primary tumor (in blue) and the brain metastases (in red), compared with data (circles). T0 = time of first cancer cell, T1 = time of diagnosis, T2 = detection time of the first brain metastasis. Only brain metastases that will become visible are shown. B. Predicted size distribution of the brain metastases at diagnosis. C. Predicted size distribution of the brain metastases at the time of clinical occurrence of the first one. *Simulations performed with the discrete version of the model. PT = primary tumor, BM = brain metastasis. Time is in months from diagnosis*.

### Generalization to a second patient

To test whether the dormancy model was generalizable, we used data from a second patient, which was not employed during the model development phase. Given the different histological type of the lung PT (squamous cell carcinoma), we adapted the doubling time accordingly (Table S1) and found a younger age of the PT of 2.1 years. Estimation of the log-kill parameter from the TGI PT model during therapy suffered from lack of identifiability, due to an estimated short duration of treatment effectiveness (see parameter *κ* in Table 2). Response of the PT to therapy was indeed characterized by a faster relapse (parameter *α*_1_), compared with the first patient. The qualitative structure of the model confirmed its descriptive power by being able to give an accurate description the BMs size dynamics (Figures S7-8), while the “basic” and “secondary dissemination” models were not (data not shown). Several parameters appeared to be patient-dependent, such as the dormancy duration *τ*, estimated to 171 days versus 133 days for the first patient. Due to the lower number of data points available for this patient, parameters identifiability was found worse, but still acceptable (Table 2).

Resulting clinical predictions were distinct (Figure S8), emphasizing the patient-specific nature of BM dynamics. The first BM was clinically detected 14 months after diagnosis, but was predicted to have been disseminated 45 months prior to diagnosis. While for both patients cerebral dissemination had already occurred at the time of diagnosis, its extent was different with a much larger mass (>1,000 cells) for the first patient than for the second (8 cells). This is due to the long period of dormancy for patient 2, resulting in all 8 predicted BMs being still dormant at the time of diagnosis.

These results demonstrate the flexibility of our model, which is able to describe and predict distinct situations in terms of repartition of the metastatic burden into individual lesions.

## Discussion

Using both population- and individual-level data of BM development in NSCLC, we have developed a general method based on biologically grounded computational models that allowed: 1) to infer the disease age from data on PT size and histology only, 2) to test multiple scenarios of metastatic dissemination and colonization against macroscopic data available in the clinic and 3) to infer times of BM initiations and number and size of invisible lesions during the clinical course of the disease.

Estimation of the duration of the pre-diagnosis PT growth phase has important implications, since BMs are more likely to have occurred for an old PT compared to a young one. Our results showed a significant difference whether considering exponential or Gompertz growth, which was consistent with previous findings^48^. Of note, our age estimates of 5.3 and 2.1 years-old are in relative agreement with the 3-4 years range found by others relying on a different method^51^.

Importantly, we were then able to include this description of PT past growth into a mechanistic model to describe the probability of BM occurrence. We found that the cell-scale dissemination parameter *μ* was significantly larger in adenocarcinomas than in squamous cell carcinomas, which offers a quantitative theory for the reported differences in BM occurrence between these two histological types^50^.

More generally, our computational platform provided a way to translate biological findings into clinically useful numerical tools. However, in order to provide robust inference, the complexity of the models had to remain commensurate with the available data. Thus, several higher order phenomena relevant to the metastatic process were ignored or aggregated into mesoscopic parameters. For instance, the metastatic potential *μ* is the product of several cell-scale probabilities relating to the multiple steps of the metastatic cascade^56^. The median values inferred from population analyses based on probabilities of BM (*μ*_pop_ = 1.39 × 10^−11^ and 1.76 × 10^−13)^ matched relatively well the ones that were found from analysis of patient-specific data (*μ* = 2 × 10^−12^ and 1.02 × 10^−12^), given the standard deviations. The larger value found in patient 1 could be due to histology or to the *EGFR* mutational status, known to impact on the BM aggressiveness^3,22,57,58^. Interestingly, while relying on distinct modeling techniques, the value of *μ* that we obtained was in the same range as obtained by others using stochastic evolutionary modeling^34,59^.

When evaluating the descriptive power of multiple biologically-based models against longitudinal data of BM sizes, our results suggested possible phases of dormancy. This finding is supported by several preclinical and clinical reports^13,60^. For instance, Nguyen et al. mention possible dormancy phases for cerebral metastases from lung cancer^13^. Durations of these latent periods are mentioned to be of the order of months, which is consistent with our quantitative results (4.4 and 5.7 months for patients 1 and 2, respectively). BM dormancy could possibly be explained by the fact that colonizing lung carcinoma cells face a foreign micro-environment when reaching the brain, which hampers their establishment and growth^61^. This could cause single cells to remain quiescent or tumors to remain at a small, pre-angiogenic size^12,62^. Indeed, vascular endothelial growth factor-A (*VEGF*-A) has been shown to mediate BM dormancy in a mouse model of lung cancer^60^.

Several second-order phenomena were ignored here for the sake of identifiability, but could nevertheless impact BM dynamics and dormancy. These include tumor-tumor interactions, either through soluble circulating factors^63^ distantly inhibiting growth and possibly maintaining dormancy^62^, or by exchange of tumor cells between established lesions^9,64^. We have recently proposed a model for distant interactions that was validated in a two-tumors experimental system and could be incorporated into the current modeling platform^65,66^.

While not problematic for the two patients that were investigated here since BMs did not respond to the systemic treatment – possibly because of the blood-brain barrier hampering delivery of the anti-cancer agents – a major limitation of our model is that the effect of therapy on the BMs was neglected. We intend to address this in future work, in particular for optimizing and personalizing combination therapies^24,67,68^. Moreover, the current study needs to be extended to a larger number of patients.

Our study demonstrates the potential clinical utility of our computational platform as a personalized predictive tool of BM that could be helpful in the decision of WBRT versus localized treatment only. For instance for patient 1, at the time of detection of the first BM or at 40 months when only 6 BMs were visible, large amounts of invisible BMs were predicted by the model, which would advise in favor of WBRT. In contrast, a smaller metastatic burden was predicted in patient 2, thus advising for localized intervention only. Thus, our quantitative modeling approach could complement existing predictive metrics such as the brain metastasis velocity index^69,70^, for patients undergoing stereotactic radiosurgery (SRS), i.e. patients presenting with a small (≤ 3-5) number of BMs. While the latter requires data at a second time point after a first SRS, our model could predict the invisible burden of BM already at the first SRS, and consequently inform the risk and timing of future relapse.

In order to translate our findings into a clinically usable tool, further methods need to be developed to calibrate the parameters of the model from data already available at diagnosis or at the first BM occurrence. By relying on a small number (3) of free parameters for the BM dynamics, our model represents a realistic candidate for such a purpose. To this respect, in addition to routine clinico-demographic features, molecular gene expression signatures^71^ (for parameter *μ*, for instance) as well as radiomics features^72^ (for parameter *γ*) might represent valuable resources.

## Materials and methods

### Patient data

The data used in this study concerned patients with non-small cell lung cancer (NSCLC) and were of two distinct natures: 1) population data of probability of BM as a function of PT size retrieved from the literature^50^ and 2) longitudinal measurements of PT and BM diameters in two patients with NSCLC retrieved from imaging data (CT scans for lung lesions, MRI for brain tumors). Both patients had unresectable PT at diagnosis. The first patient (used for model development) was extracted from an EGFR mutated cohort from Institut Bergonié (Bordeaux, France). The second patient (used for model validation) had an EGFR wild type squamous cell lung carcinoma and was issued from routinely treated patients in the thoracic oncology service of the University Hospital of Marseille. The data comprised 10 PT sizes and 47 BM sizes (spanning 6 time points) for the first patient and 11 PT sizes and 16 BM sizes (spanning 4 time points) for second patient.

Data use from the Marseille patient was approved by a national ethics committee (CEPRO, Comité d’Evaluation des Protocoles de Recherche Observationnelle, reference number 2015-041), according to French law. Data use from the Marseille patient was approved by the institutional ethics committee of Institut Bergonié (Comité de Protection des Personnes Sud-Ouest et Outre Mer III) and IRB approval was obtained for use of the images. The need for written informed consent of data collection was waived for this patient, in accordance with the related policy of Institut Bergonié. The study was performed in accordance with the Declaration of Helsinki, Good Clinical Practices, and local ethical and legal requirements.

### Mathematical modeling of primary tumor growth and metastatic development

#### Primary tumor growth

The pre-diagnosis natural history of the primary tumor size *S*_p_(*t*) for times *t* < *T*_d_ (diagnosis time) – expressed in number of cells in the model computations – was assumed to follow the Gompertz growth model^46,47^, i.e.

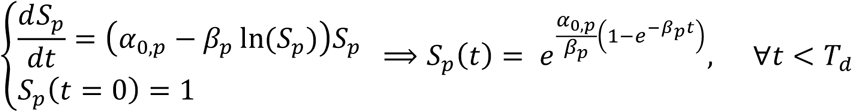

where time *t* = 0 corresponds to the first cancer cell, parameter *α*_0,p_ is the specific growth rate 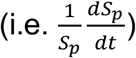 at this time and *β*_p_ is the exponential rate of decrease of the specific growth rate. Conversions from diameter measurements to number of cells were performed assuming spherical shape and the classical assumption 1 mm^3^ = 10^6^ cells^73^. After treatment start (*T*_d_), the primary tumor size was assumed to follow a tumor growth inhibition model^54^ consisting of: 1) exponential growth (rate *α*_1_), 2) log-kill effect of the therapy (efficacy parameter *κ*)^74^ and 3) exponential decrease of the treatment effect due to resistance, with half-life *t*_res_. The equation is:

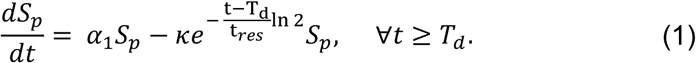

#### Metastatic development

##### (A, B). Basic and different growth models

The general modeling framework we employed was derived from the work of Iwata et al.^31^. It consists in modeling the population of metastases by means of a size-structured density *ρ*(*t*, *s*), of use to distinguish between visible and invisible tumors. Metastatic development of the disease is reduced to two main phases: dissemination and colonization^75^. The multiple steps of the metastatic cascade^56^ are aggregated into a dissemination rate with expression:

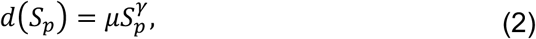

which corresponds to the number of successfully born BM per unit of time. In this expression, the geometric parameter *γ* corresponds to the intra tumor repartition of the cells susceptible to yield metastasis and *μ* is the per day per cell probability for such cells to overcome all the steps of the metastatic cascade (acquisition of metastatic-specific mutations, epithelial-to-mesenchymal transition, invasion of the surrounding tissue, intravasation, survival in transit, extravasation and survival in the brain). For *γ* = 1 all cells in the PT have equal probability to give a BM whereas a value of *γ* = 0 indicates a constant pool of cells having metastatic ability (cancer stem cells). Intermediate values 0 < *γ* < 1 can be interpreted as the geometric disposition of the metastatic-able cells, including the surface of the tumor (*γ* = 2/3) or a fractional dimension linked to the fractal nature of the tumor vasculature^76^. Assuming further that the growth of the metastasis follows a gompertzian growth rate

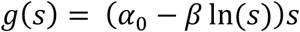

with growth parameters *α*_0_ and *β* possibly equal (basic model (A)) or distinct (different growth model (B)) compared to the PT ones, the density ρ satisfies the following transport partial differential equation^31^:

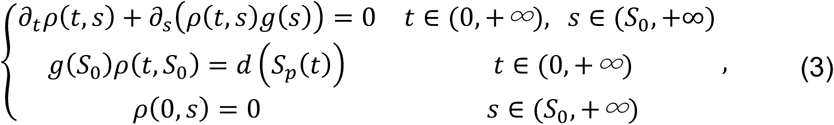

where *S*_0_ is the size of a BM at birth (here assumed to be one cell). From the solution to this equation, the main quantity of interest for comparison to the empirical data is the number of metastasis larger than a given size s (cumulative size distribution):

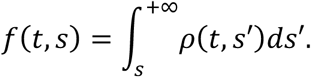

The total number of metastases – denoted *N*(*t*) – is obtained by using *s* = *V*_0_ above and its expression can be directly computed without solving the entire problem (3) as it is given by:

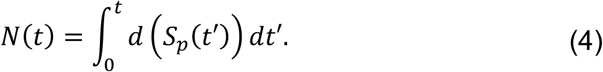

Using the method of characteristics, one can derive the following relationship between *N* and *f*:

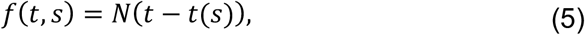

where *t*(*s*) is the time for a tumor growing at rate *g* to reach the size s. In the case of Gompertz growth one has:

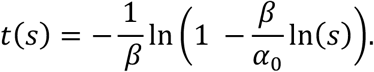

Of particular interest is the number of visible BMs *f*(*t*, *S*_vis_) with *S*_vis_ the minimal visible size at CT scan taken here to be 5 mm in diameter.

##### (C). Secondary dissemination

In the previous model formulation, all BMs were assumed to have been seeded by the primary tumor. When BMs are also allowed to spread metastases themselves, this results in a second term in the boundary condition of (3) and the model becomes^31^:

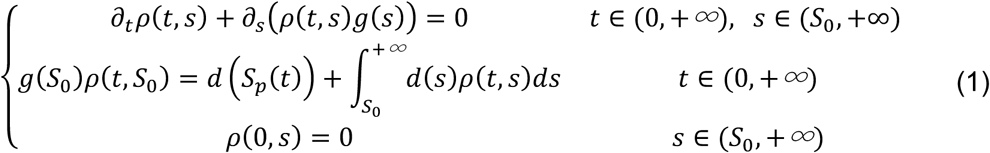

In this case, formula (4) is not valid anymore, which complicates substantially the computation of the cumulative size distribution. A dedicated scheme based on the method of characteristics was employed^43^.

##### (D). Delay model

Consideration of a delay *t*_0_ before onset of metastatic dissemination in the model can be taken into account by remarking that

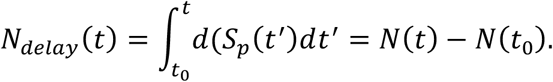

Thus

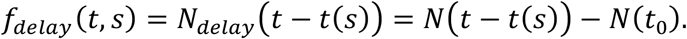

##### (E). Dormancy model

For inclusion of dormancy in the model – defined as a period of duration τ during which a newborn metastasis remains at size *S*_0_ – the time to reach any given size *s* > *S*_0_ becomes *t*_dorm_(*s*) = *t*(*s*) + *τ*. The cumulative size distribution is then given by:

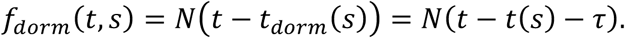

##### Discrete versions of the models

While continuous versions of the models were used for fitting the model to the data because they allow computations to be tractable, discrete versions were implemented for forward calculations, because of the small number of BMs. Briefly, in a discrete simulation, the appearance time of the *i* -th BM *T*_i_ was defined by

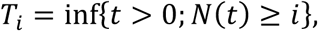

with *N*(*t*) the number of metastases as defined above. The size of the *i*-th BM *S*_*i*_(*t*) was then defined, for *t* > *T*_i_, by:

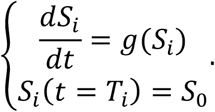

This discrete version corresponds to a Poisson process where the number of events (metastatic births) *N*(*t*) is replaced by its expectation: *N*(*t*) = E[*N*(*t*)]. For further links between stochastic (Poisson process) and continuous versions of the Iwata model, the reader is referred to^77^. Simulations reported in Figures 5B-C and 6 were generated using discrete versions of the models.

### Models fitting and estimation of the parameters

#### Pre-clinical primary tumor growth

To parameterize the Gompertz function defining the PT growth, two parameters needed to be defined (*α*_0,*p*_ and *β*_*p*_). In the absence of longitudinal measurements of the PT size without treatment, these two parameters were determined from two considerations: 1) the maximal reachable size (carrying capacity, equal to 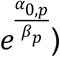 of a human tumor is 10^12^ cells^29,49^ and 2) the histology-dependent value of the doubling time at diagnosis, retrieved from a meta-analysis of published literature about the natural growth of lung PTs (see Table S1, extended from^44,48^). The latter yielded values of 201 days for an adenocarcinoma and 104 days for an undifferentiated carcinoma. For the Gompertz model, the doubling time is size-dependent and its value for the PT diagnosis size *DT*(*S*_*d*_) is given by:

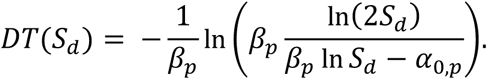

Using the formula linking *α*_0,*p*_ and *β* to the carrying capacity, this nonlinear equation was numerically solved.

#### Population level: probability of BM apparition

To fit the data from^50^ describing the probability of BM in a population of lung adenocarcinoma patients, we employed a previously described methodology^39,78^. Briefly, we considered that the probability of developing BM after diagnosis was the probability of having already BM at diagnosis, i.e. P(*N*(T_*d*_) > 1), with *N* defined by equation (4). Of note, since this quantity only depends on the dissemination rate, this probability is equal for all the model scenarios introduced above. We fixed the value of the PT growth parameters as described above from the cohort histology and set *γ* = 1 as the simplest dissemination model. The inter-individual variability was then minimally modeled as resulting from a lognormal population distribution of the parameter *μ* (ln *μ* ∼ *N*(ln(*μ*_*pop*_), *μ_σ_*)). Uniform distributions of the PT diameters were assuming within each interval (*S*^*i*^, *S*^*i*+1^) of the PT sizes S^1^, …, *S*^6^given as data. The probability of developing a metastasis with a PT size *S* ∈ (*S*^*i*^, *S*^*i*+1^) writes:

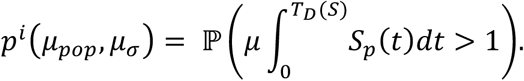

The best-fit of these probabilities – evaluated by Monte Carlo simulations – to the empirical data was then determined by least squares minimization performed using the function *fminsearch* of Matlab (Nelder-Mead algorithm)^79^.

#### Individual level: description of longitudinal data of number and size of BM growth

##### Maximum likelihood estimation

Due to the discrete nature of the data at the individual level (diameters of a small number of BMs at discrete time points), a direct comparison between the size distribution *ρ* solution of the problem (3) was not possible. Instead, we compared the data to the model by means of the cumulative size distribution. Denoting by *t*_*j*_ the observation times, *x*_*j*_^*i*^ the sorted BM sizes at time *t*_*j*_ and y_*j*_^*i*^ the number of metastases larger than size *x*_*j*_^*i*^ at time *t*_*j*_, we considered the following nonlinear regression problem:

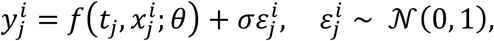

where *θ* = (*α*_0,*p*_, *β*_p_, *α*_0_, *β*, *μ*, *γ*, *t*_0_, *τ*) regroups all the parameters of the model. Note that, except for the “secondary dissemination” model, all models can be viewed as submodels of a general model including all these parameters (the “basic model” consisting of the case *α*_0,*p*_ = *α*_0_, *β*_*p*_ = *β*, *t*_0_ = 0 and *τ* = 0, for instance). Classical maximum likelihood estimation then leads to the following estimate:

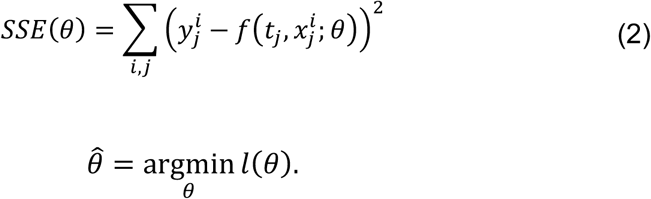

##### Parameters identifiability

Standard errors can be computed from this statistics’ covariance matrix, given by^80^:

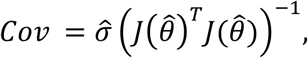

where *J*(*θ̂*) is the jacobian matrix of the model (with respect to the parameter vector *θ*) at all time points and all sizes, evaluated at the optimal parameter *θ̂* and *σ̂* is the a posteriori estimate of *σ* given by

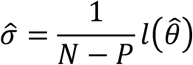

with *N* the total number of data points and *P* the number of free parameters.

Using the standard errors as an identifiability metric, we repeatedly observed a lack of identifiability of parameters *μ* and *γ* when fitted together. Indeed, standard errors for *μ* and *γ* were larger than 200% when fitting the basic model to the data. Further investigation of the shape of the objective function confirmed this lack of idenfiability (Figure S7). To address this issue, we only considered a finite set of relevant possible values for *γ* and only optimized the value of *μ*. These values were (0, 0.4, 0.5, 2/3, 1) and corresponding initial conditions for *μ* were (10^-3^, 10^-5^, 10^-8^, 10^-9^, 10^-12^). When more parameters were let free in the model (delay *t*_d_ or dormancy period *τ*), we generated 4 × 4 parameters grids for initial conditions (with *t*_d_ ∈ {0,500,1700,2000} or *τ* ∈ {0, 50, 100, 350}). Of these 16 optimization problems, the one with the minimal value of *l* at convergence was selected.

## Acknowledgments

The authors would like to thank Pr A. Iliadis for valuable comments and advice.

## Author contributions

SB and DB conceived the research idea. CS, AB, PT, FB and FC collected the data. CP measured tumor sizes. MB and SB performed the data analysis. MB and SB programmed the software for estimation of the parameters and simulation. The paper was written by SB with editorial input from all authors.

The authors declare no potential conflicts of interest

## Data availability statement

The datasets analyzed during the current study are available from the corresponding author on reasonable request

## Supplementary Material

**Figure S1:**
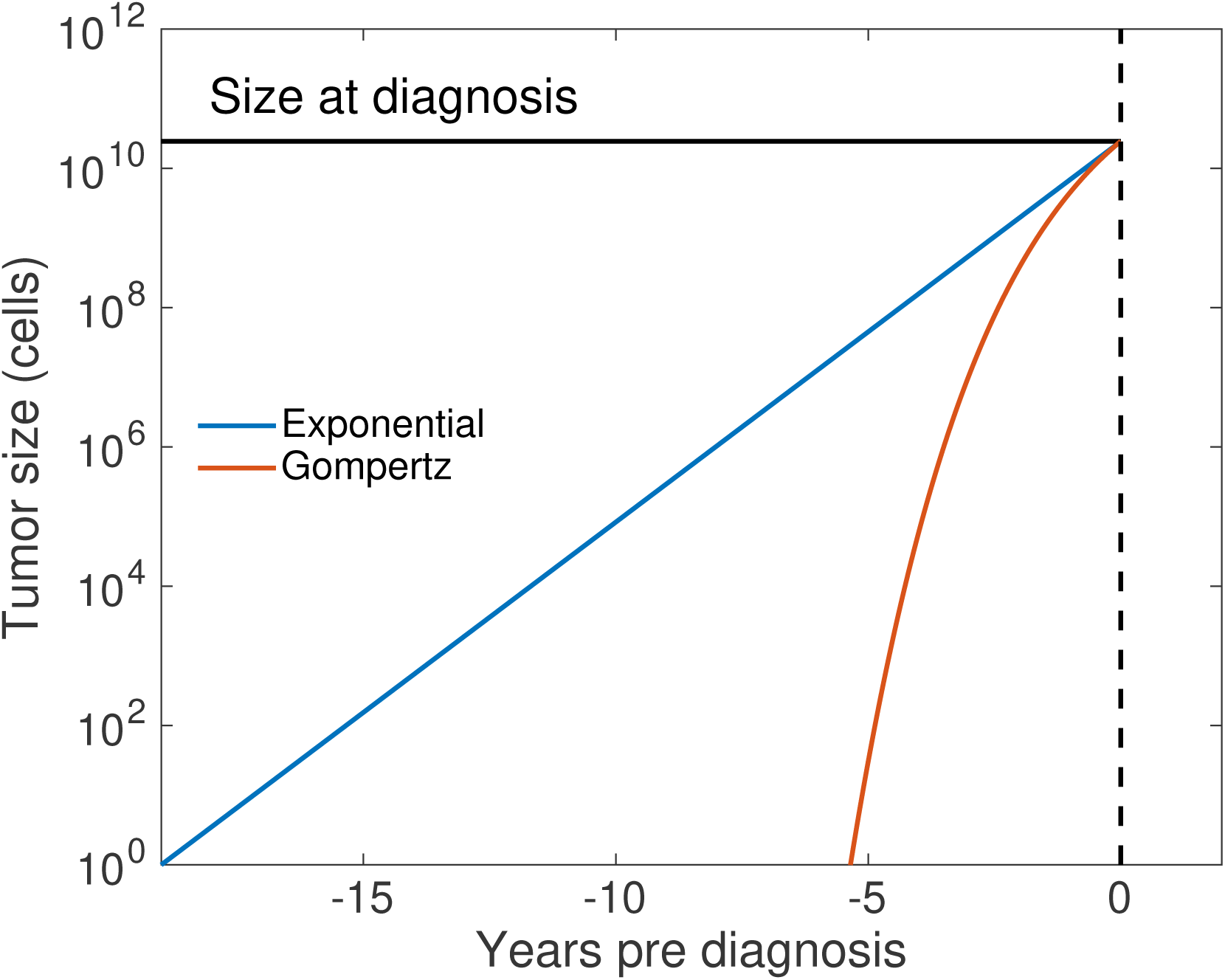
Comparison of exponential and Gompertz predictions of pre-clinical phase of growth

**Figure S2:**
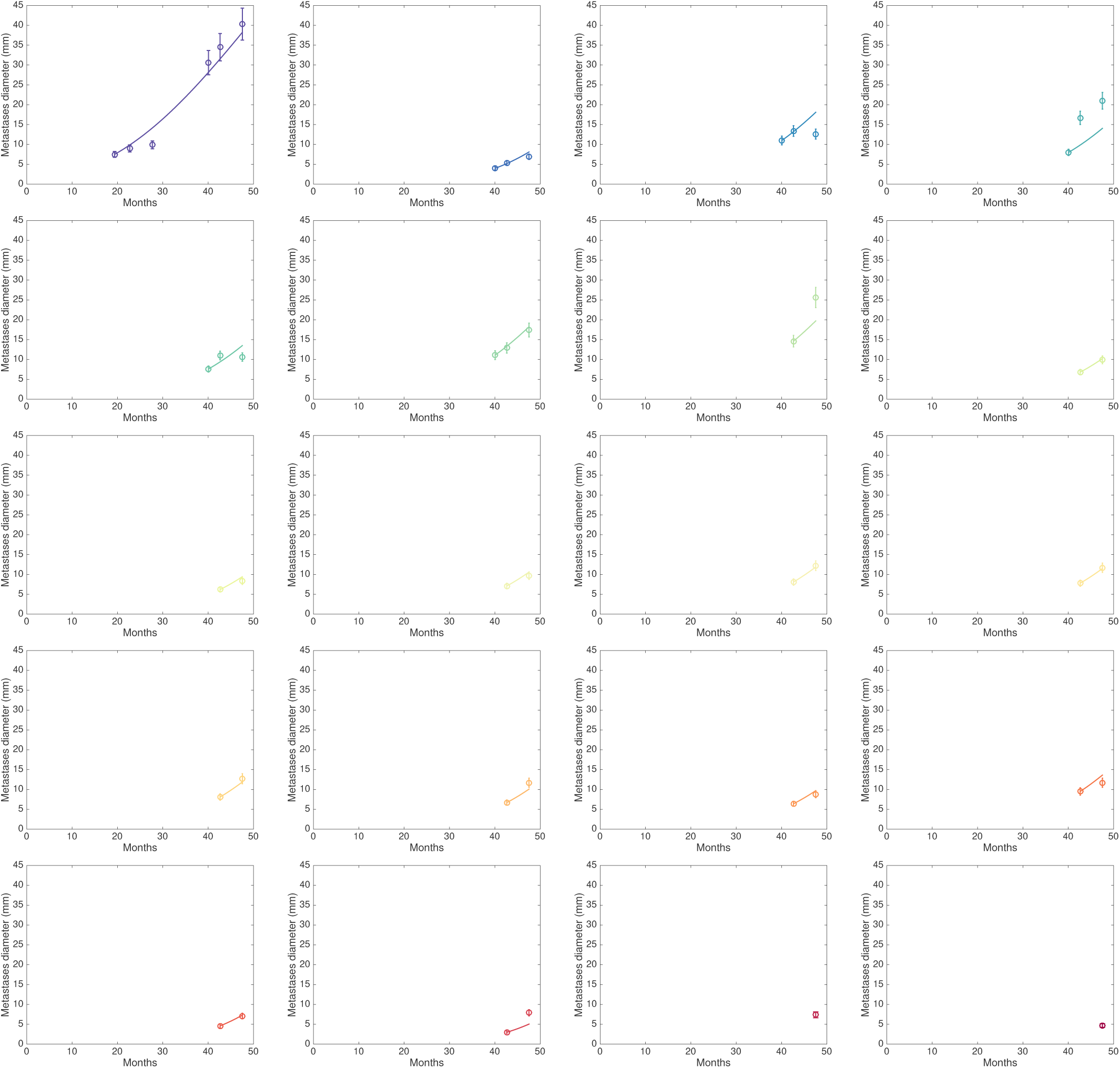
Gompertz growth predictions of individual brain metastases. Comparison between the Gompertz growth model predictions and the lesion size data of individual BMs. The Gompertz growth parameters were determined only from the primary tumor size at diagnosis and histology (adenocarcinoma, see Methods). Apart from the first measurements used as initial conditions, no other future measurements were used to make the predictions. Error bars represent 10% error on the size measurement.

**Figure S3:**
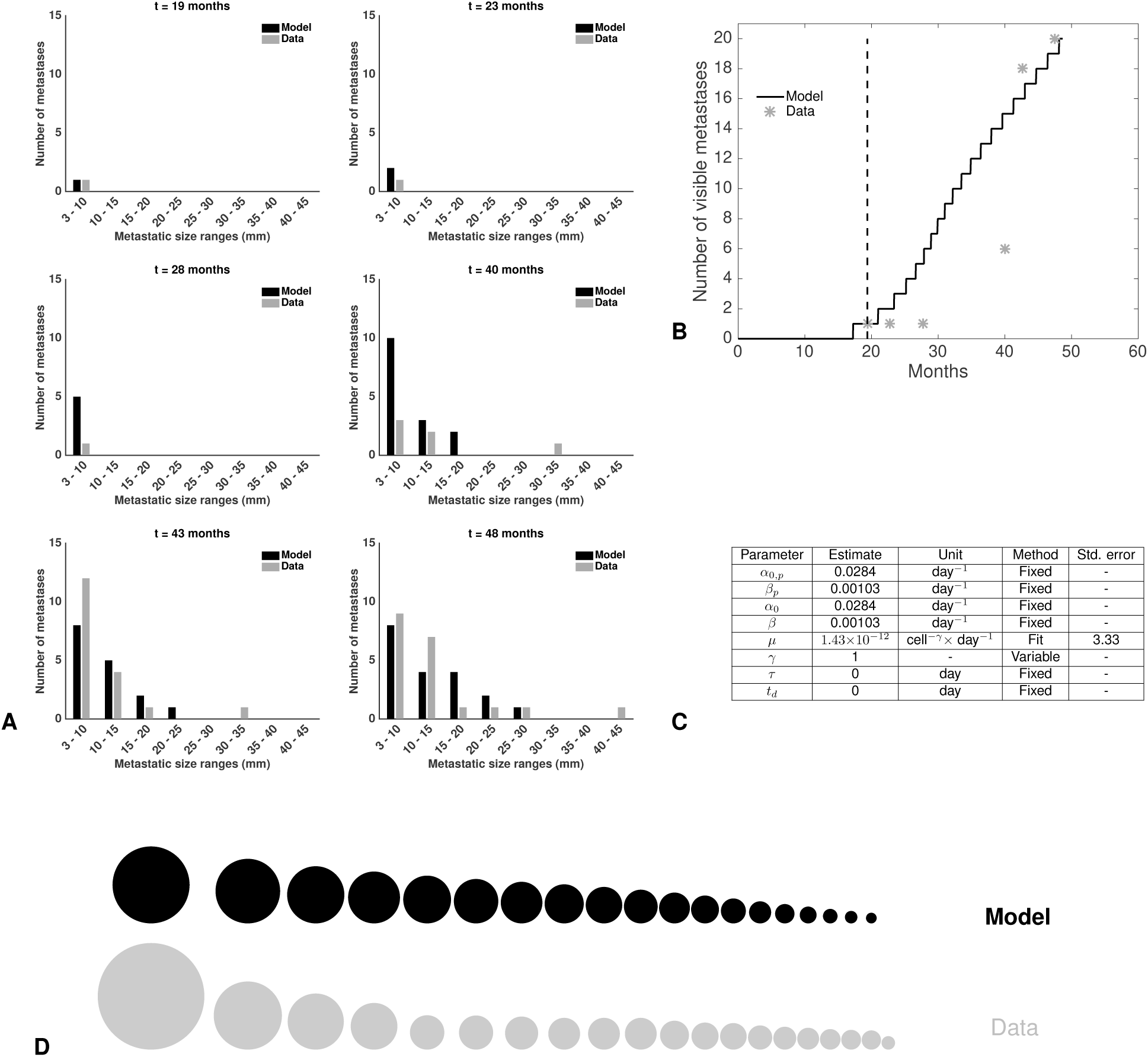
Fit of the basic model. A. Time course of the visible brain metastases (BMs) size distributions during follow-up. Comparison between model calibration and data. B. Time course of the number of visible BMs. C. Parameter estimates. Std error = Standard errors expressed in percent. D. Comparison of the BM size distribution between the model fit and the data at the last time point.

**Figure S4:**
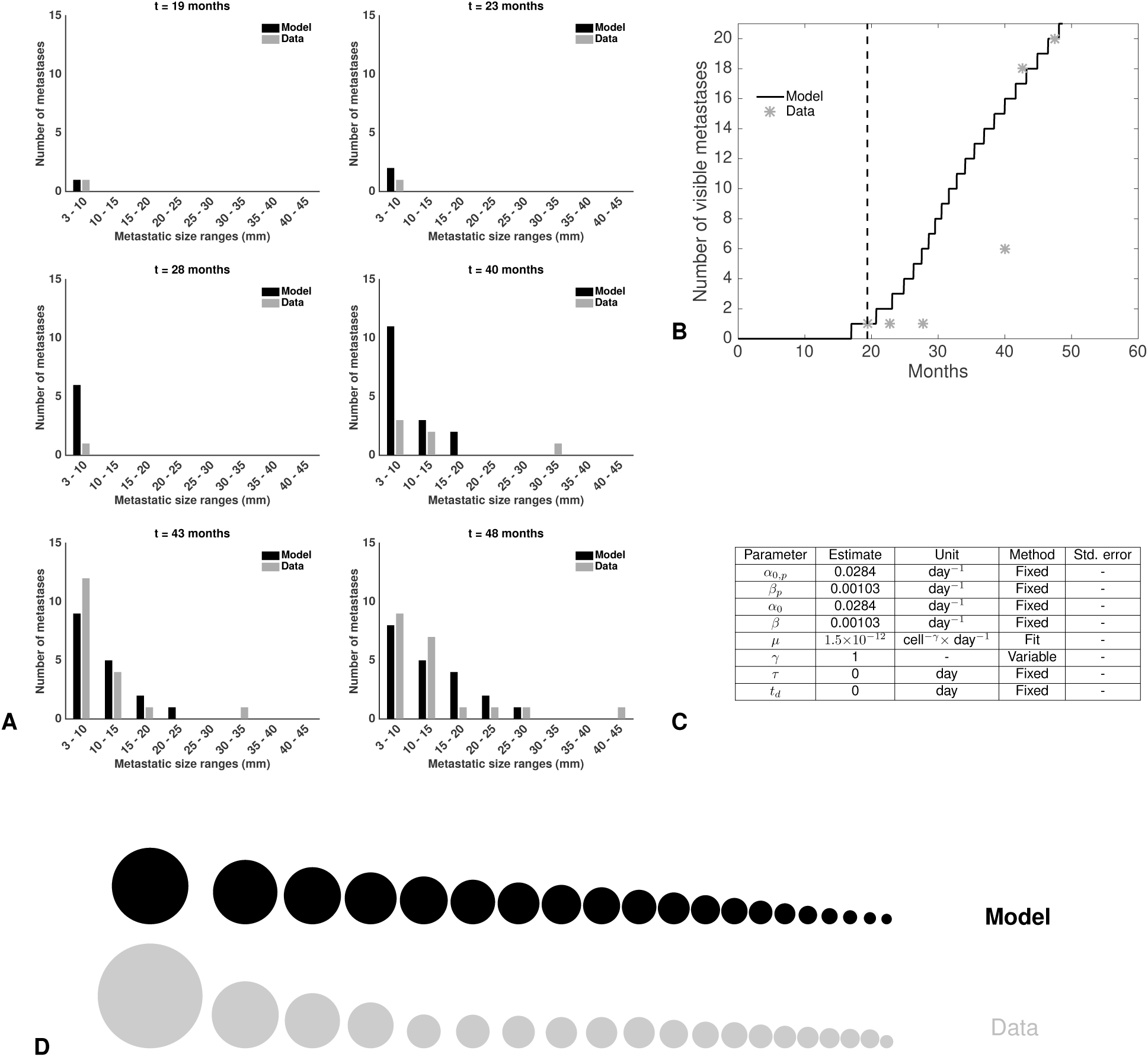
Fit of the model with secondary dissemination. A. Time course of the visible brain metastases (BMs) size distributions during follow-up. Comparison between model calibration and data. B. Time course of the number of visible BMs. C. Parameter estimates. Std error = Standard errors expressed in percent. D. Comparison of the BM size distribution between the model fit and the data at the last time point.

**Figure S5:**
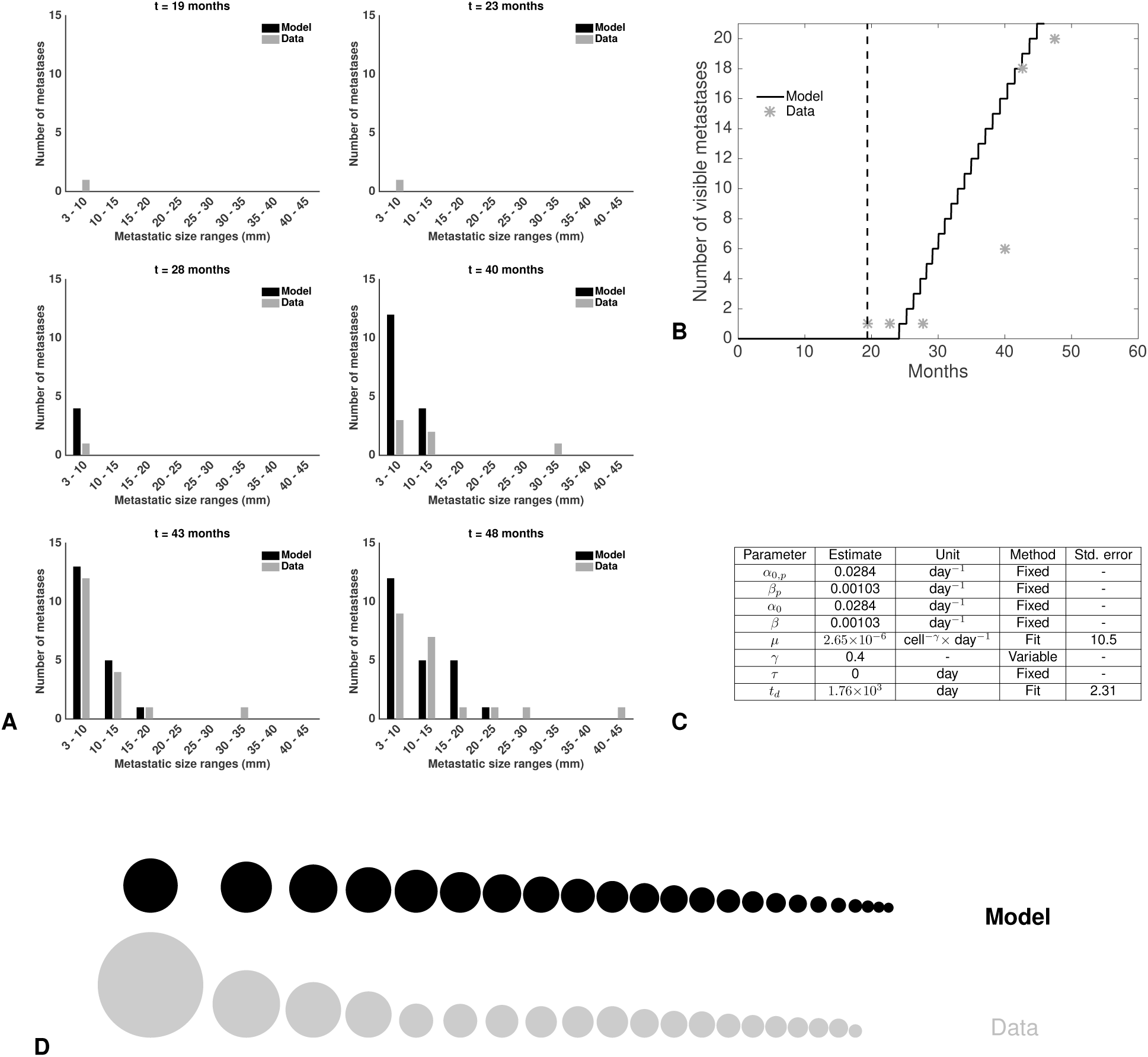
Fit of the delay model. A. Time course of the visible brain metastases (BMs) size distributions during follow-up. Comparison between model calibration and data. B. Time course of the number of visible BMs. C. Parameter estimates. Std error = Standard errors expressed in percent. D. Comparison of the BM size distribution between the model fit and the data at the last time point.

**Figure S6:**
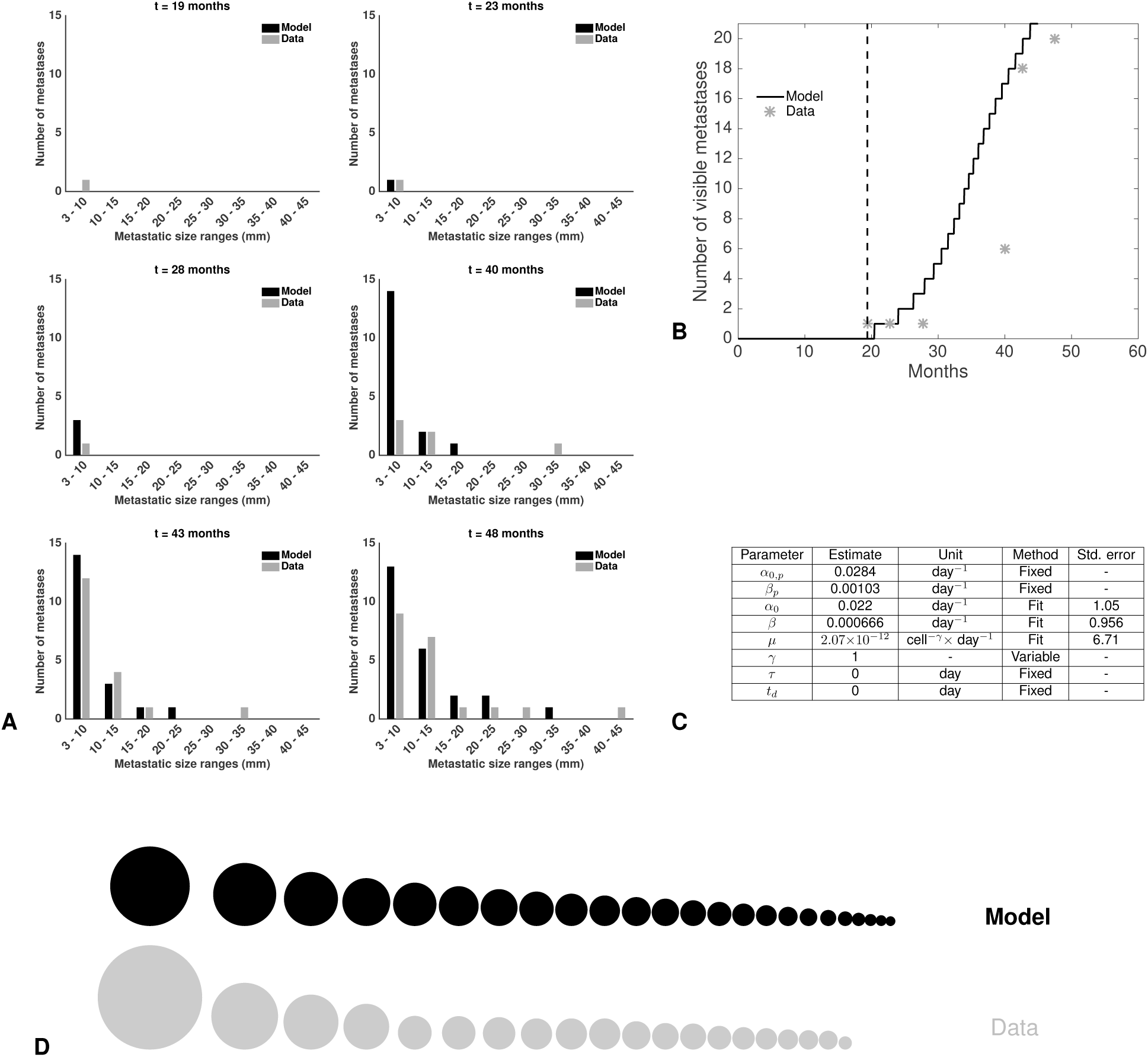
Fit of the model with different primary and secondary growth parameters. A. Time course of the visible brain metastases (BMs) size distributions during follow-up. Comparison between model calibration and data. B. Time course of the number of visible BMs. C. Parameter estimates. Std error = Standard errors expressed in percent. D. Comparison of the BM size distribution between the model fit and the data at the last time point.

**Figure S7:**
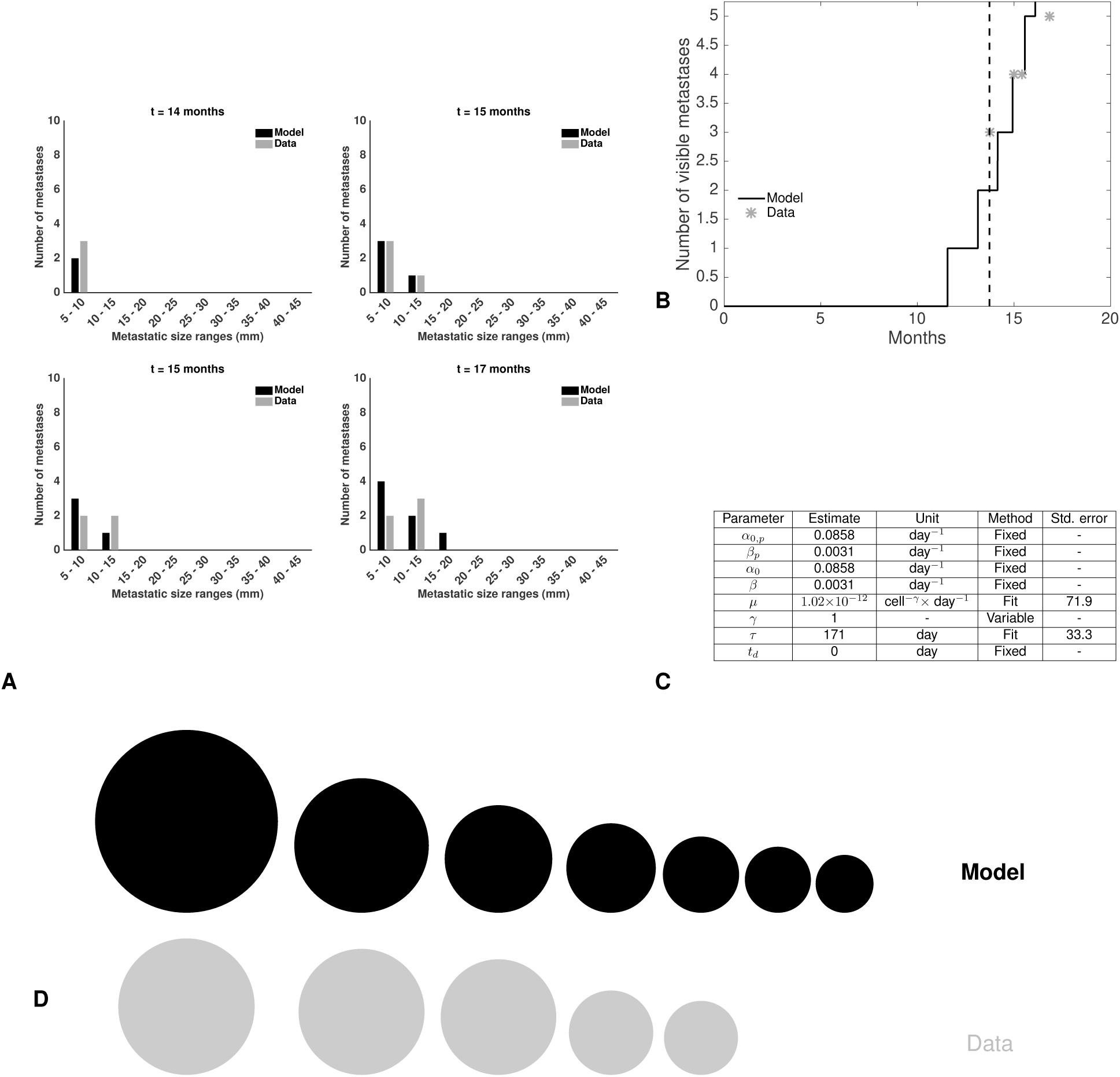
Fit of the dormancy model for patient 2. A. Time course of the visible brain metastases (BMs) size distributions during follow-up. Comparison between model calibration and data. B. Time course of the number of visible BMs. C. Parameter estimates. Std error = Standard errors expressed in percent. D. Comparison of the BM size distribution between the model fit and the data at the last time point.

**Figure S8:**
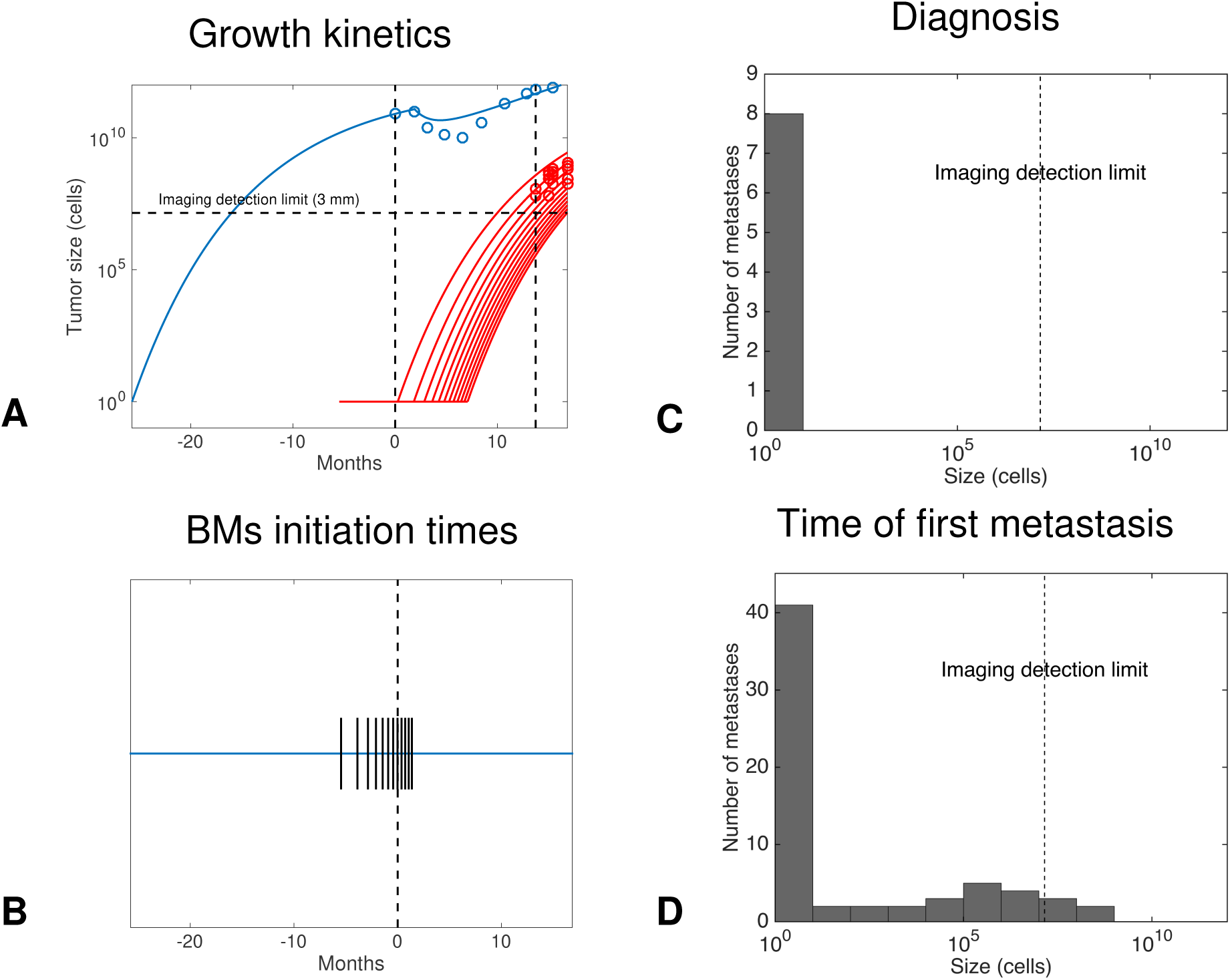
Predictions for patient 2. A. Inferred growth kinetics of the primary tumor (in blue) and the brain metastases (in red). Only brain metastases that will become visible are shown. B. Model predictions of the initiation times of the brain metastases. C. Predicted size distribution of the brain metastases at diagnosis. D. Predicted size distribution of the brain metastases at the time of clinical occurrence of the first one.

**Figure S9:**
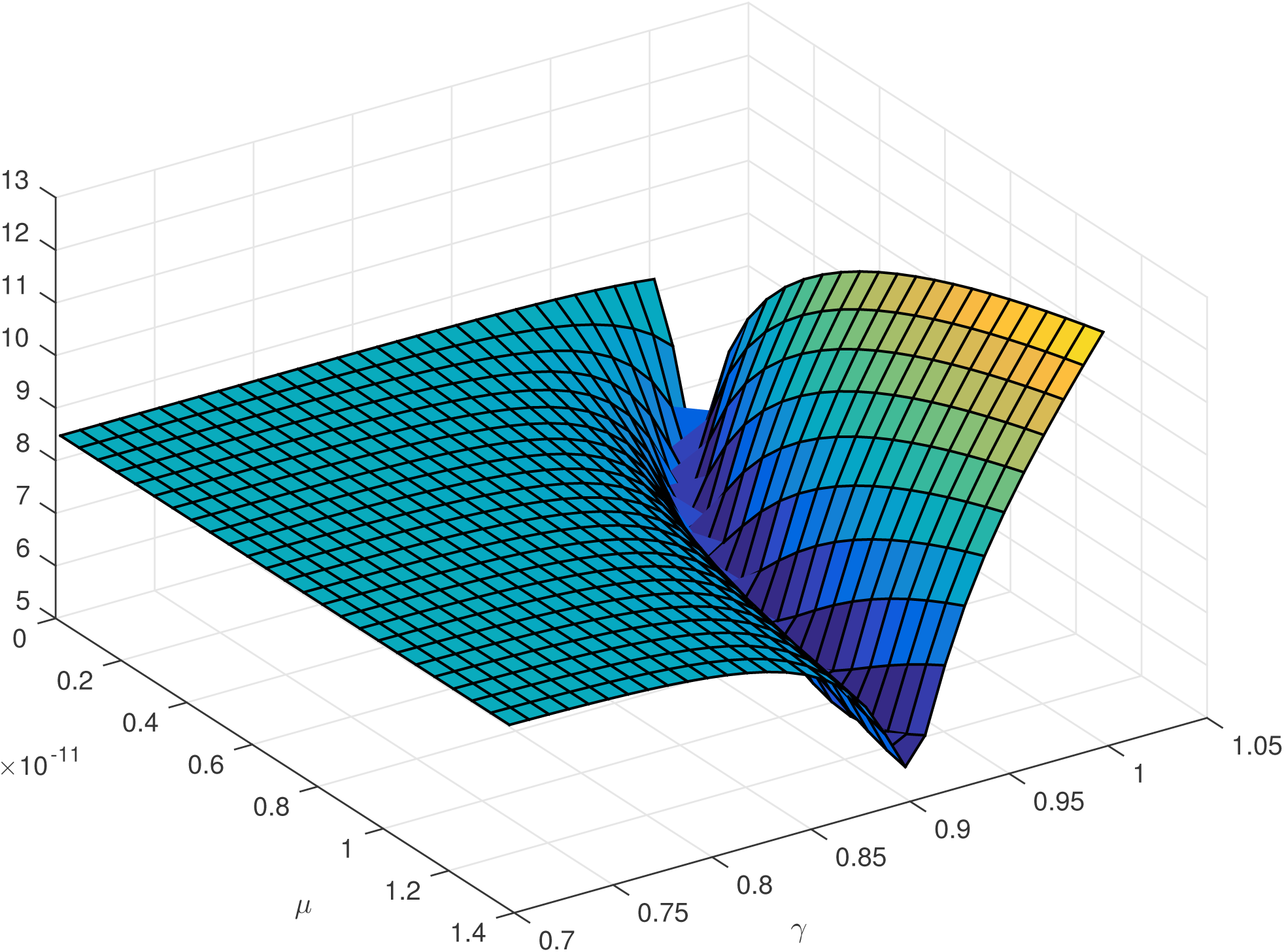
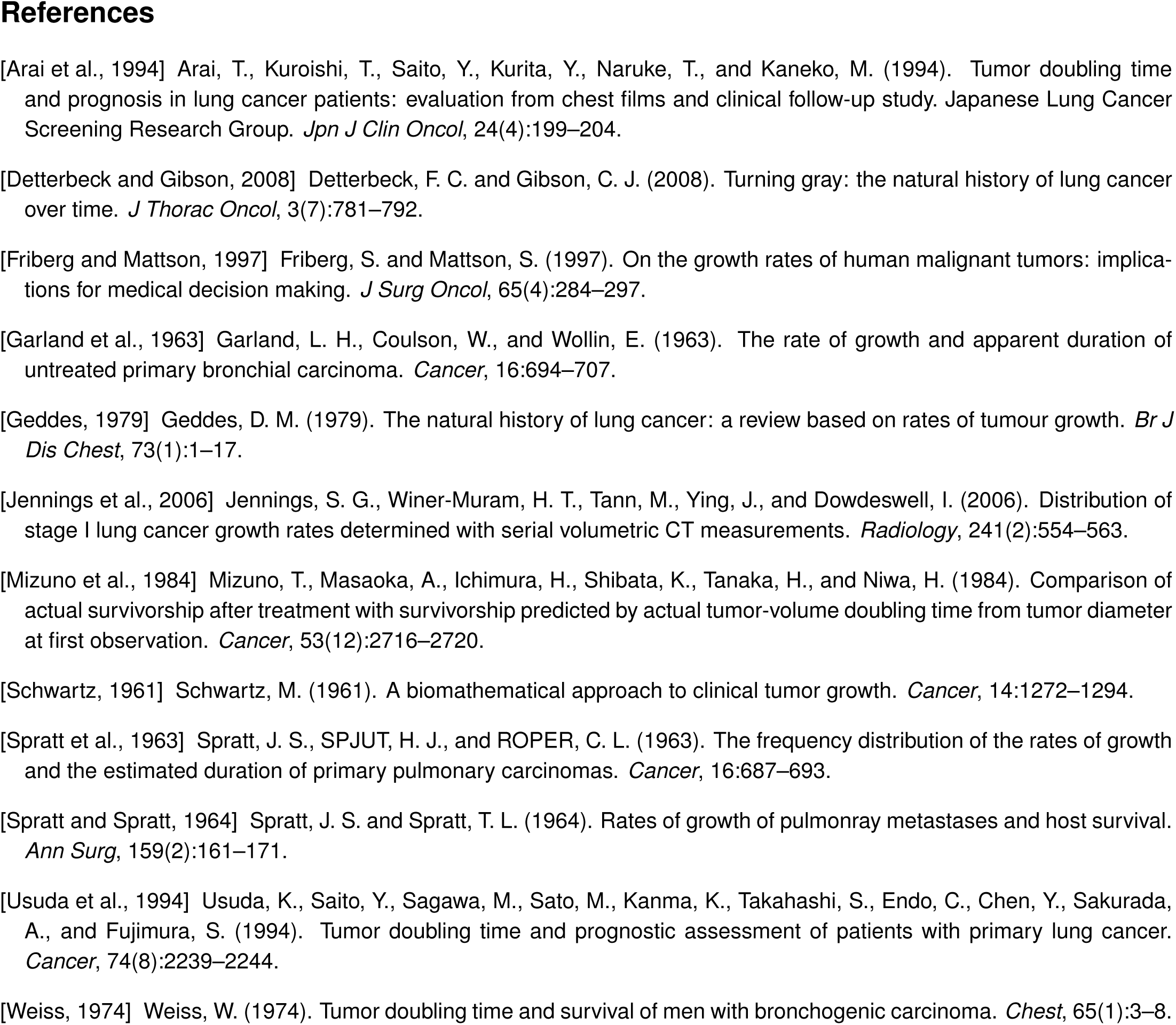
Shape of the objective as a function of *µ* and *γ*.

**Table S1:** Growth kinetics of primary lung tumors

| Ref. | All | Adenocarcinoma | Squamous cell carcinoma | Undiff. |
| --- | --- | --- | --- | --- |
| [Jennings et al., 2006] | 161 (149) | 166 (51) | 132 (48) |  |
| [Usuda et al., 1994] | 164 ± 178 (165) | 223 ± 209 (86) | 105 ± 106 (67) | 79 ± 52 (12) |
| [Arai et al., 1994] | 166 (237) | 222 (133) | 115 (69) | 68 (9) |
| [Mizuno et al., 1984] | 136 (50) | 178 ± 157 (23) | 103 ± 78 (22) | 111 ± 57 (5) |
| [Geddes, 1979] | 102 (228) | 161 (60) | 88 (111) | 86 (44) |
| [Weiss, 1974] | 183 ± 85 (28) | 214 ± 73 (11) | 78 ± 29 (8) | 109 ± 72 (7) |
| [Spratt and Spratt, 1964] | 88 (34) | 118 (8) | 70 (13) | 93 (13) |
| [Spratt et al., 1963] | 112 (22) | 269 (7) | 93 (6) | 90 (9) |
| [Garland et al., 1963] | 162 (41) | 222 (7) | 126 (22) | 123 (9) |
| [Schwartz, 1961] | 78 ± 48 (13) | 72 ± 77 (2) | 79 ± 46 (11) |  |
| **Average\*** | **143** | **201** | **104** | **91** |

Reports of volume doubling times from routinely detected primary NSCLC tumors from the literature (extended from [Detterbeck and Gibson, 2008] and [Friberg and Mattson, 1997]). Note: squamous cell carcinoma = epidermoid carcinoma and undifferentiated carcinoma cell carcinoma = large cell carcinoma. All denotes all histology from lung cancer together (including both NSCLC and small cell carcinomas).
Undiff. = undifferentiated carcinoma.
Mean ± standard deviation.
In parenthesis are the number of patients.
\*Average was computed as the weighted mean of each study with weights the number of patients.

## Notes

#### Summary of Updates

Substantial revision of Figures and Text to improve clarity.

